# Classifying cold stress responses of inbred maize seedlings using RGB imaging

**DOI:** 10.1101/432039

**Authors:** Tara A. Enders, Susan St. Dennis, Justin Oakland, Steven T. Callen, Malia A. Gehan, Nathan D. Miller, Edgar P. Spalding, Nathan M. Springer, Cory D. Hirsch

## Abstract

Increasing the tolerance of maize seedlings to low temperature episodes could mitigate the effects of increasing climate variability on yield. To aid progress toward this goal, we established a growth chamber-based system for subjecting seedlings of 40 maize inbred genotypes to a defined, temporary cold stress while collecting digital profile images over a 9-day time course. Image analysis performed with PlantCV software quantified shoot height, shoot area, 14 other morphological traits, and necrosis identified by color analysis. Hierarchical clustering of changes in growth rates of morphological traits and quantification of leaf necrosis over two time intervals resulted in three clusters of genotypes, which are characterized by unique responses to cold stress. For any given genotype, the set of traits with similar growth rates is unique. However, the patterns among traits are different between genotypes. Cold sensitivity was not correlated with the latitude where the inbred varieties were released suggesting potential further improvement for this trait. This work will serve as the basis for future experiments investigating the genetic basis of recovery to cold stress in maize seedlings.

## Introduction

Climate change threatens to negatively impact performance of many important crops, including maize. Extreme heat and drought in maize can cause decreases in yield, especially during later stages of development (Sánchez et al., 2014). One method of avoiding yield losses due to extreme heat and drought late in the season is to plant crops earlier in the season (Kucharik, 2008); however, earlier planting increases the risk of exposing maize seedlings to low temperature stress conditions.

Cold stress is often described as a freezing stress (≤ 0°C) or a chilling stress (generally above 0°C and below 15°C) across plant species (Lyons, 1973; Greaves, 1996). Suboptimal temperatures can have multiple impacts on plant growth depending on the severity and developmental time point at which the stress occurs. Effects can range from slight delays in development from growth inhibition to plant death. Other commonly observed stress responses include leaf chlorosis and necrotic lesions (Yadav, 2010).

As a species, maize is considered cold-sensitive (Sellschop and Salmon, 1928); however, genetic variation in cold sensitivity exists among inbreds (Greaves, 1996). Several studies have considered maize genotypes that display mild cold sensitive phenotypes to be cold tolerant, despite the lines still being affected by cold stress (Janowiak and Dörffling, 1996; Fracheboud et al., 1999; Sowiński et al., 2005; Wijewardana et al., 2015). However, it is difficult to try to compare levels of sensitivity across studies done under different growth conditions, different temperatures, and at different developmental stages. Also, previous studies have rarely analyzed more than two maize genotypes at a time. Greaves (1996) stated that to improve plant performance under low temperature conditions, genetic variation needed to be characterized for multiple traits, such as levels of tissue injury and growth rates. To identify optimal genetic material for breeding programs interested in maximizing cold tolerance in maize, it is essential to thoroughly characterize the range of cold sensitivity.

Many physiological processes in plants are impeded by low temperatures, such as photosynthetic capacity, membrane rigidity, transpiration, and enzyme activity (Marocco et al., 2005). Together, these physiological effects of cold stress can result in poor agronomic performance, such as slower emergence, decreased biomass accumulation, reduced growth rates, and leaf chlorosis and necrosis (Miedema, 1982). Relative growth rates (Hetherington and Oquist, 1988; Verheul et al., 1996), electrolyte leakage assays (Capell and Dörffling, 1993), and photosynthesis related measurements (Hetherington and Oquist, 1988; Aguilera et al., 1999) have been used in several studies to classify maize seedling responses to cold stress. These approaches require destructive measurements of plants, and therefore necessitate more individuals and space to collect time course data. However, some measurements have been conducted in a non-destructive manner. In field conditions, necrotic injury was visually assessed on a relative scale at a single time point on six genotypes, where lines with the least amount of leaf necrosis were classified as cold tolerant (Janowiak et al., 2003). An alternative approach to destructive and manual assessment is to move to image-based plant phenotyping methods to allow for robust measures of changes in color and growth without destructive, subjective, or labor-intensive techniques.

There are a growing number of commercial and custom-built systems for integrated controlled plant growth and imaging (Tisné et al., 2013). Currently available commercial systems can provide valuable insight into variation in plant growth and development, but they tend to have higher costs and infrastructure requirements that limit access to a small number of researchers. Additionally, these systems usually restrict researchers to conducting a single experiment at a time. We sought to develop an image-based approach that could be implemented easily to document morphological traits of maize seedlings at a low cost. This system is not fully automated because it requires manual plant staging. As manual staging of plants is required, we have implemented tools to ensure high-quality standardized images are captured.

Many researchers have, or are currently developing, custom low-cost phenotyping platforms to fit their needs. Our system is not necessarily unique in this pursuit and is similar to other recently developed low-cost imaging systems (Knecht et al., 2016; Armoniené et al., 2018; Czedik-Eysenberg et al., 2018). Currently, our system is limited to acquiring images of seedlings from one side view image but has the advantage of being scalable. Additionally, as this system was developed separately from a greenhouse or growth chamber, it can be used to image concurrently running experiments.

Numerous software tools to analyze plant traits from images are available (www.plant-image-analysis.org; Lobet, 2017). The ImageJ plugin HTPheno measures height, width, and area from side-view plant images (Hartmann et al., 2011). Integrated Analysis Platform contains pipelines for multiple plant species and can output measurements such as height, width, skeleton length, volume, convex hull, and number of leaves (Klukas et al., 2014). ImageHarvest measures multiple traits, such as plant dimensions, shoot area, convex hull area, and center of mass (Knecht et al., 2016). PlantCV has functions for multiple shape measurements, such as height, width, perimeter length, center of mass coordinates, and others (Fahlgren et al., 2015; Gehan et al., 2017). The recent addition of a Naive Bayes classifier to PlantCV allows users to quantify color-based features of plants (Gehan et al., 2017). We chose to use PlantCV for plant trait extraction, so that, in addition to morphological features, we could easily quantify leaf necrosis effects of cold stress manifesting as color changes in leaf tissue in our image-based data set.

This study sought to compare growth rates of 40 diverse maize inbreds in a mild cold stress treatment under controlled growth conditions using image-based phenotyping methods. We established an image acquisition platform, including a system for embedding metadata and sample tracking, to collect high-quality, standardized RGB images of maize seedlings over time. Trait extraction from images was accomplished using PlantCV. This work describes a robust method for analyzing recovery rates across multiple morphological and color-based traits for a large number of maize inbreds that will serve as a foundation for future work uncovering the genetic basis for cold stress recovery in maize.

## Methods

### Plant Material and Growth Conditions

Forty genotypes were used in this study, including: 3IIH6, B73, CM105, CM37, CML052, CML069, CML103, CML228, CML277, D06, DK105, F2, F353, F7, HP301, Il14H, Ki11, Ky21, LH185, LH198, LH82, M162W, M37W, Mo17, Mo18W, MoG, MS71, NC350, NC358, Oh43, Oh7B, P39, PH207, PHJ89, PHP02, Tx303, Tzi8, UH007, W117, and W22. Details for each genotype, such as developer, market class, and population group, are provided in Supplemental Table 1. For all experiments, seeds were planted in 40 cubic inch D-40 DeePots (Stuewe and Sons, Inc.) containing a 1:1 mix of SunGro (Agawam, MA) horticulture professional growing mix and autoclaved field soil approximately 2 inches below the surface. The plants were grown in Conviron growth chambers with a 16 hour 30°C and 8 hour 20°C day/night cycle and watered every other day. The cold stress treatments were implemented using a Thermo Scientific refrigerated incubator programmed with a 16 hour 6°C and 8 hour 2°C day/night cycle. Plants were moved to cold stress conditions approximately 2 hours after dawn at 9 days after sowing (DAS) for the indicated amount of time as required for various experiments. After the indicated time of stress treatment, treated plants were moved from the cold incubator back to growth chambers under control temperature conditions.

### Image Acquisition

A Nikon D5100 DSLR Camera with an 18-55mm lens mounted on a Provista 7518B Tripod (Davis & Sanford) produced digital RAW-format images (Figure 1A). A computer running the Ubuntu 14.04 operating system was interfaced with the camera through its universal serial bus. A single shell script that combined gPhoto2 (http://gphoto.org), dcraw https://www.cybercom.net/~dcoffin/dcraw/, tiffcp (http://www.libtiff.org), and MatLab functions controlled image acquisition, assessed image quality, and converted each RAW image file into tagged image file format (TIFF). Custom MatLab code checked that the appropriate camera settings for focal length, f-number, exposure, and body tilt matched defined values. If an image failed the quality and standardization checks, the script identified the problem and prompted the user to retake the image. Approved images in RAW format were automatically stored in a directory corresponding to the date of image acquisition. Sample tracking information and experimental metadata were encoded in a two-dimensional barcode (Quick Response code format) and printed on a piece of paper that was mounted in the scene above the plants (Figure 1A). This embedded experimental details and plant identity into the corresponding image data in a machine-readable format. The sample tracking page also contained 24 blue boxes. The user marked the number of blue boxes corresponding to the age of the plants each day they were photographed. This date/age score was automatically added to the sample tracking data at the time of image acquisition.

**Figure 1.**
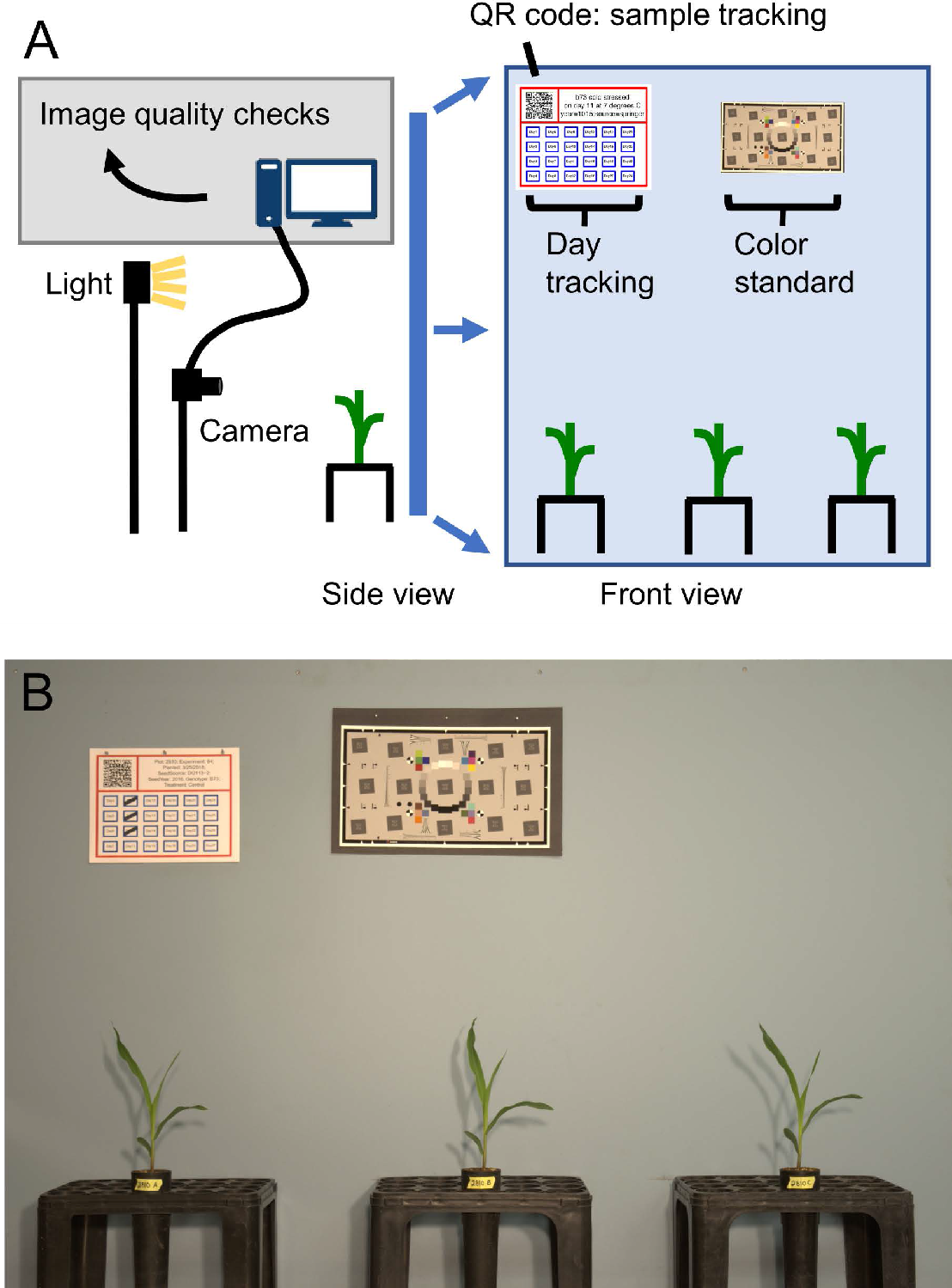
Schematic of the RGB imaging system. A) The imaging system consisted of a DSLR camera, computer, back lighting, blue background, and three racks to hold pots. Metadata for each image was stored in a QR code and day after sowing was crossed off on a sheet of paper hung on the top left of the background in each image. A color standard was also included for use in image analysis pipelines. B) Representative example of an acquired image.

We refer to the three seedlings in each image as a plot. The staging area consisted of a desk, a 4′x6′ blue drywall background, three plastic D20T racks (Stuewe and Sons, Inc), nails to hold the QR code and day tracking sheets, and the space to the right of the QR code was used to add color standards but could be used for other information as well. The use of DeePots provided several advantages in the system as plants could be grown at high densities and easily moved from growth chambers to the imaging system. Additionally, DeePots enabled plants to be quickly moved in and out of racks, rotated as necessary to adapt to rotations in growth, and placed in consistent locations each day. An example image acquired using this system is depicted in Figure 1B. This system can be used for imaging a wide variety of plant species to obtain side views of plants over time. The combination of imaging platform size and growth conditions allowed for growth and for data to be collected from 8 to 16 days after sowing (DAS). After this time, the growth of the seedlings began to plateau in some genotypes and the leaves of neighboring plants began to overlap, which hindered proper plant segmentation.

### Generation of Sample Metadata Tracking Sheets

Custom Perl and R scripts were written to create QR code sheets containing metadata and to allow sample tracking over time. Briefly, a Perl script is run that takes in a tab delimited text file containing the desired sample tracking metadata (plot, genotype, treatment, etc). This Perl script outputs an R script that can be run to produce the formatted QR code sheet with embedded metadata and day tracking boxes. These scripts are available at https://github.com/maizeumn/cold-phenotyping.

### Trait extraction from images using PlantCV

Raw .nef format RGB files of maize seedlings were converted to .tiff format files using dcraw (https://www.cybercom.net/~dcoffin/dcraw/). Trait measurements were extracted from each .tiff file using PlantCV v3.0.dev2 (Fahlgren et al., 2015; Gehan et al., 2017; doi:10.5281/zenodo.1408271). Pixel classification within each image was achieved through the use of the Naive Bayes multiclass training module within PlantCV. This approach allowed for color-based classification of plant tissue into two categories: healthy and necrotic, and therefore quantification of the percent area (number of pixels) corresponding to each of these categories (Figure 2A). Plant masks were dilated and filled to reduce noise in the segmentation. Initial efforts to classify plant pixels in this manner resulted in soil pixels being included in the plant necrotic category. Ranges of RGB values for necrotic tissue, stem tissue, and soil overlapped and could not be separated using our training set and the Naive Bayes approach. To remove soil pixels from the plant mask, we identified the rack that held each pot using edge detection methods and excluded any pixels below a boundary line to isolate only plant pixels for later trait extraction. The final plant mask and original RGB image were used to measure attributes of the plant object.

**Figure 2.**
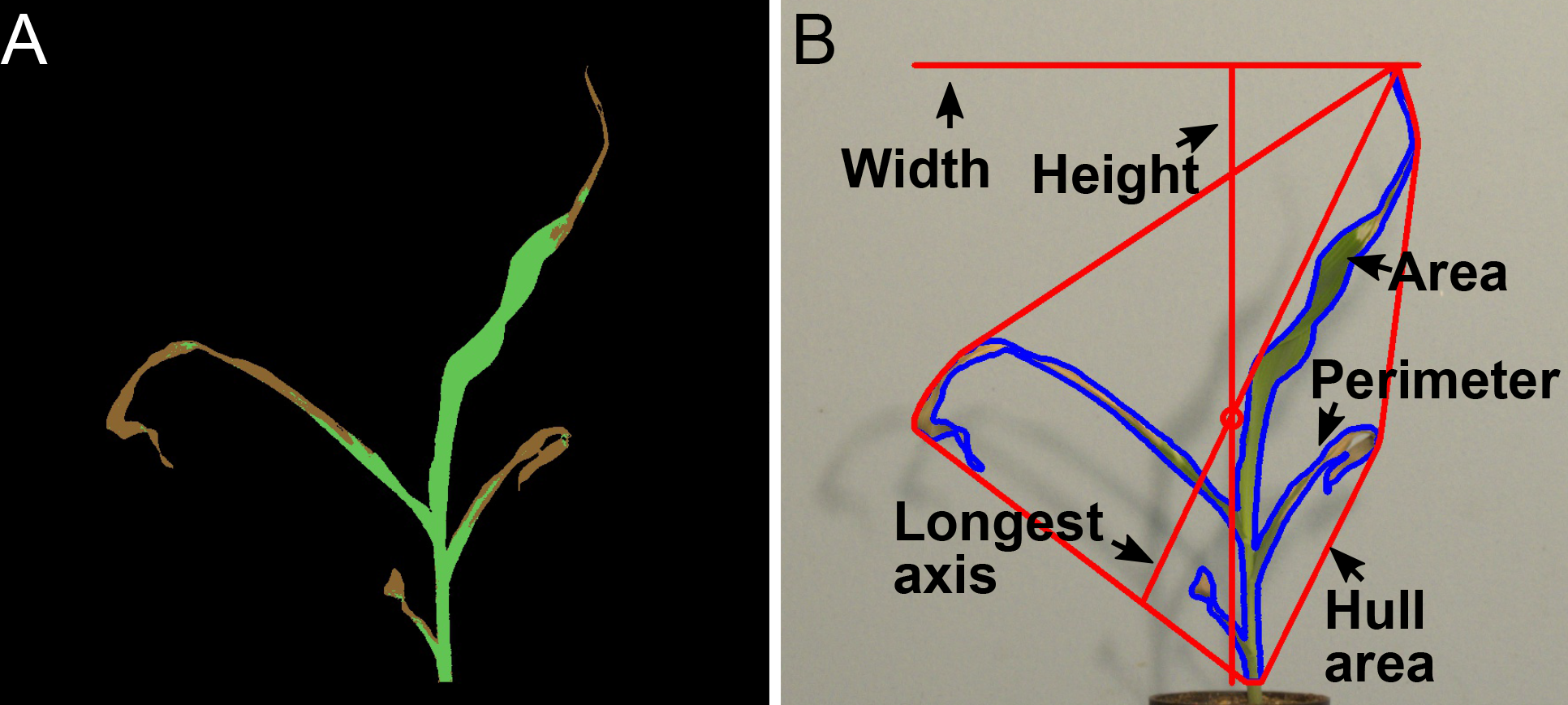
Extracted traits. A) False-colored image indicating the quantification of the areas classified as necrotic (brown) and healthy (green) tissue. B) Plant with overlaid representative measured morphological traits.

Our pipeline used PlantCV to measure 16 morphological traits associated with defined objects, such as height, width, area, convex-hull properties, and various measurements of an object-bounding ellipse (Figure 2B). Height represented the number of pixels from the base of the stem to the tallest point of the plant object. Width captured the number of pixels along a horizontal line between the plant pixel with the smallest x-coordinate to the highest x-coordinate. Area was defined as the number of pixels classified as plant tissue, including pixels in both healthy and necrotic categories. Perimeter represented the number of pixels along the outermost edge of the plant object. Measurements derived from the convex hull included the area of the convex hull, the number of vertices of the convex hull, the longest axis within the convex hull, and solidity. Ellipse-derived measurements included x and y coordinates of the ellipse center, major and minor axis of the ellipse, the rotational angle, and the eccentricity. Traits that were measured by hand had high correlations to the image-derived measurements, suggesting that our image acquisition and trait extraction techniques yielded accurate quantification of these aspects of plant growth. Some morphological traits corresponded to readily explained descriptors of plant morphology, and we chose to focus on traits that were reproducible and captured unique aspects of responses to cold stress in our genotypes. Numerical measurements in pixels for tissue classification categories and morphological traits were written to a .csv file for each image. We chose to output four images for each plant analyzed, which captured various processing steps and documented the quality of plant segmentation (Supplemental Figure 1). The individual .csv output files were merged using a python script, and the data was analyzed in R. Input TIFF format images are available on Cyverse Data Commons (https://doi.org/10.7946/P2T63C). Numerical outputs from PlantCV pipeline as merged .csv files and scripts including R code used to generate figures, Perl code to generate QR code and metadata sheets, and scripts for image acquisition are available here: https://github.com/maizeumn/cold-phenotyping. README files, both on Cyverse for image data and Github for scripts, provide short explanations and usage for each file provided.

### Data analysis

#### Plant Growth Rates

For experiments examining the effect of cold stresses of different durations on plant growth, points on line plots represented the mean of six plants per genotype per treatment, and error bars represent standard error of the mean. All experiments were replicated three times with similar results.

For experiments surveying responses of genotypes under a 2 day cold stress period, line plots represented the means of n≥14 plants per genotype per treatment at each timepoint. Sample sizes for each genotype for each treatment on each day are indicated in Supplemental Table 2. Where indicated, significance between treatment groups within a genotype was determined by a one-way ANOVA and post-hoc TukeyHSD test to obtain adjusted p-values.

#### Rate calculations

The growth rate for indicated time intervals for each unique combination of genotype, treatment, and trait was calculated by finding the slope of a linear regression line for each plant. The slope values were averaged to obtain a single value for each genotype, treatment, trait, and interval group.

#### Trait clustering

The pheatmap function from the R package ‘pheatmap’ was used to create the heatmap displaying hierarchical clustering of genotypes for each trait for each interval.

## Results

### Cold stress assay development

Cold temperatures often result in slower growth and induce leaf necrosis in maize seedlings, but the severity of these effects varies among genetic backgrounds (Greaves, 1996). The goal of this study was to survey the range of cold stress effects on the growth, morphology, and leaf necrosis across various maize genotypes. The first step towards accomplishing this goal was the design of a cold-stress assay that resulted in phenotypic changes across genetic backgrounds, which also allowed for analysis of recovery within the constraints of our image acquisition system.

Our system provided an opportunity to measure plant growth and morphology for maize seedlings from 8 to 16 DAS. We conducted several experiments to identify a set of conditions that allowed analysis of variability for responses to cold stress. A variety of temperatures and stress lengths were assessed to determine appropriate cold stress conditions. While temperatures near or below 0°C provided strong stress responses, we found greater experiment-to-experiment variation and some genotype lethality at these temperatures. Therefore, we elected to use a more moderate low temperature condition (6°C day / 2°C night) that was more phenotypically consistent but resulted in more subtle phenotypes than freezing temperatures. We tested the effects of different durations of this cold stress to select a treatment regime that resulted in observable effects on measured traits but also allowed for quantification of stress recovery in the B73 and Mo17 inbreds (Figure 3). All cold-treated plants were placed in the stress condition two hours after dawn at 9 DAS. Every 24 hours a subset of plants were removed from the cold stress and returned to control conditions in growth chambers for a total of four separate durations of cold treatment ranging from 1 to 4 days. Plants grown under a single 24-hour period of cold treatment had the least amount of growth inhibition in height, area, and width compared to plants grown under control conditions for both genotypes (Figure 3). Four days of cold stress resulted in the most extreme differences in growth compared to control plants for area, height, and width measurements but only allowed for 3 time points during the recovery period. A 2- or 3-day duration of cold stress had intermediate effects to these two extremes. To collect a maximum amount of time points during recovery, a 2-day cold treatment was chosen for further experiments. Additionally, the selection of a 2-day long cold stress over a 1-day cold stress allowed for the analysis of whether any of our selected genotypes were able to grow during the cold period or if the cold stress conditions resulted in growth arrest for all tested genotypes and the collection of five time points to characterize recovery from stress conditions.

**Figure 3.**
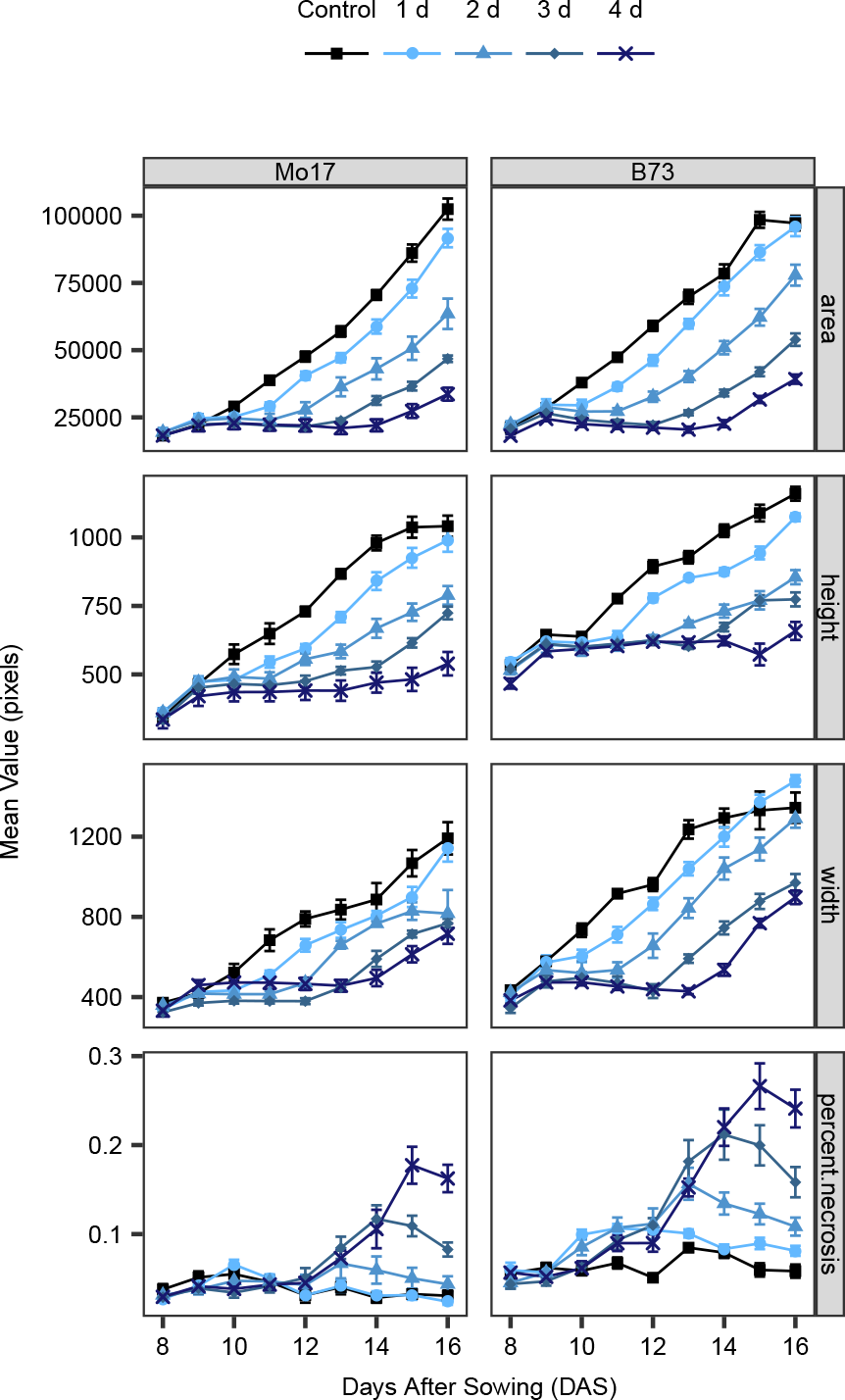
Mean growth rates of traits for B73 and Mo17 control and cold-stressed seedlings. Cold stressed seedlings were transferred to a separate incubator with a 6°C/2°C day/night cycle on day 9 after sowing, and stressed for the indicated amount of time (1, 2, 3, or 4 days), then returned to the control-temperature growth chamber. Error bars represent standard error of the mean of 6 plants.

### Surveying diversity of cold responses using image-based methods

To survey the variation in responses to cold stress among diverse genotypes of maize, we implemented a robust cold stress assay that subjected plants to two days of cold stress in growth chamber conditions to analyze cold stress responses in a panel of 40 maize genotypes. Within our selected genotypes, there were representatives from multiple heterotic groups and genotypes that resulted from breeding programs at very distinct latitudes that likely faced variable levels of early season cold stress (Figure 4A; Supplemental Table 1). The genotypes used in this study included 21 of the 25 nested association mapping population parent lines (McMullen et al., 2009), and lines with sequenced genomes such as B73 (Schnable et al., 2009; Jiao et al., 2017), PH207 (Hirsch et al., 2016), W22 (Springer et al., 2018), F7 (Unterseer et al., 2017), and Mo17 (Sun et al., 2018). All plants were grown in conditions described in the methods with a 2-day cold stress as the treatment group. This 2-day cold treatment was able to recapitulate phenotypes observed in field grown plants that experience cold stress, such as leaf necrosis, chlorosis, and growth inhibition. Additionally, our image-based data collection method enabled the capture of nine time points during the early developmental stages of maize plants (Figure 4C, D). Images of Mo17 seedlings grown under control conditions (Figure 4C) and cold-treatment conditions (Figure 4D) provide examples of the developmental time points captured for each seedling and the degree of growth inhibition achieved in our assay. We collected data on three biological replicates that represented different grow-outs. For each biological replicate, we measured traits for six individuals exposed to control conditions and six individuals exposed to a cold stress. In total, this dataset contained images for nine consecutive days of growth for ~18 plants per genotype per treatment resulting in ~12,000 images of ~1,400 plants.

**Figure 4.**
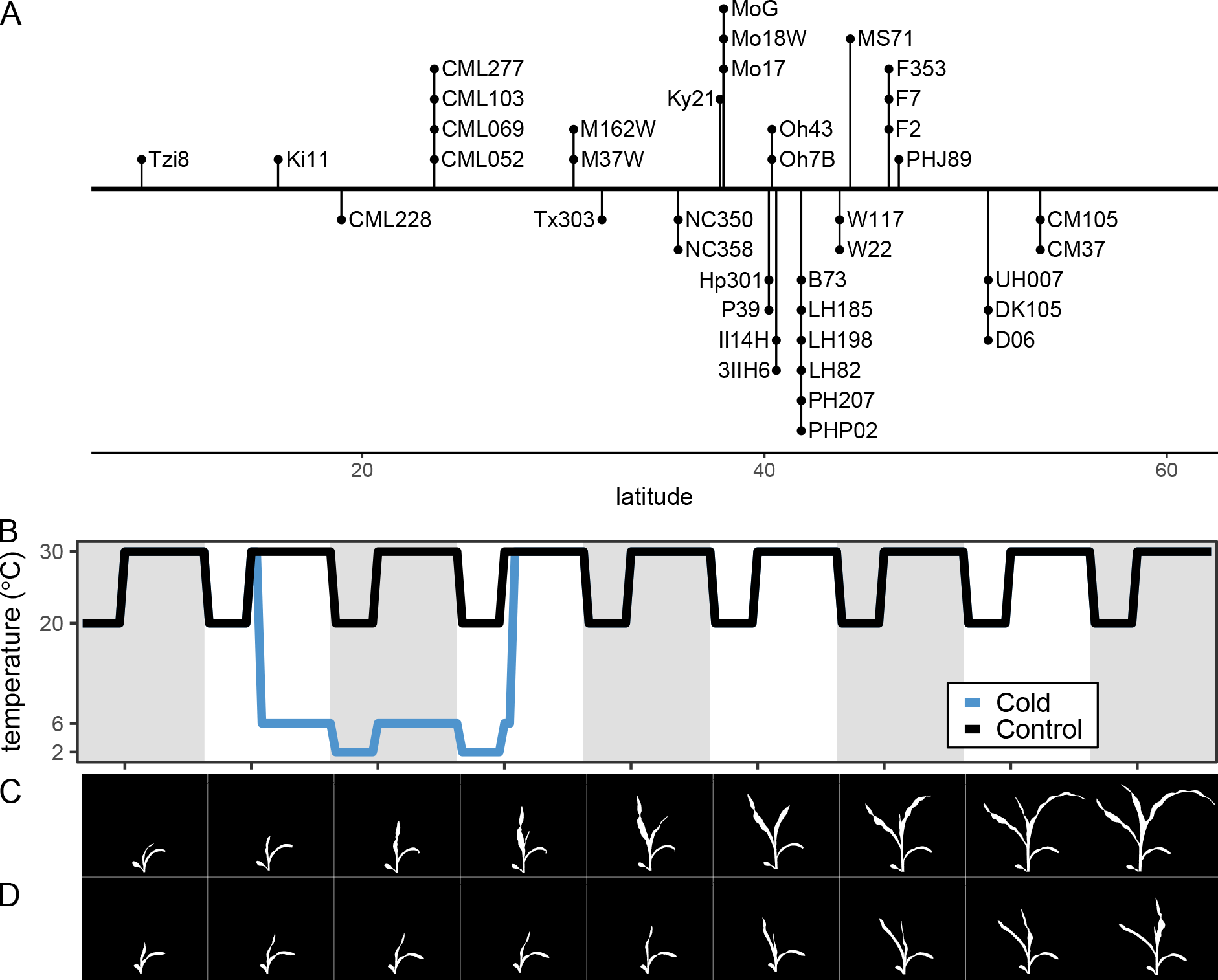
Experimental design to survey genotypic responses to cold stress. A) Latitudes of source location of the 40 inbred genotypes included in this study. B) Plants were grown under a 30°C 16-hour day and 20°C 8-hour night cycle. At 9 days after sowing, cold-stressed plants were placed in a low-temperature incubator under a 6°C 16-hour day and a 2°C 8-hour night until 11 days after sowing and returned to the control conditions. RGB images of the plants were collected between 9:00 am and 12:00 pm each day. C) Image masks of a Mo17 control plant at each time point. D) Image masks of Mo17 cold-stressed plant at each time point.

### Impact of cold stress on leaf necrosis across genotypes

One of the more noticeable effects of cold stress in maize seedlings is the appearance of leaf necrosis (Figure 5). Cold temperatures can cause leaf tissue to wilt, and over the course of several days this wilted tissue can die, resulting in changes in leaf color from green to brown and texture from healthy leaves with high turgor to dehydrated, dead leaf tissue (Guye et al., 1987). The Naive Bayes color-based classification module that was trained and implemented within PlantCV classified each plant pixel into a healthy or necrotic category that enabled the quantification of necrosis as a percentage of area belonging to each category for every plant at each time point. The pipeline output included images of each seedling indicating the category each pixel was classified into by color (Figure 2A).

**Figure 5.**
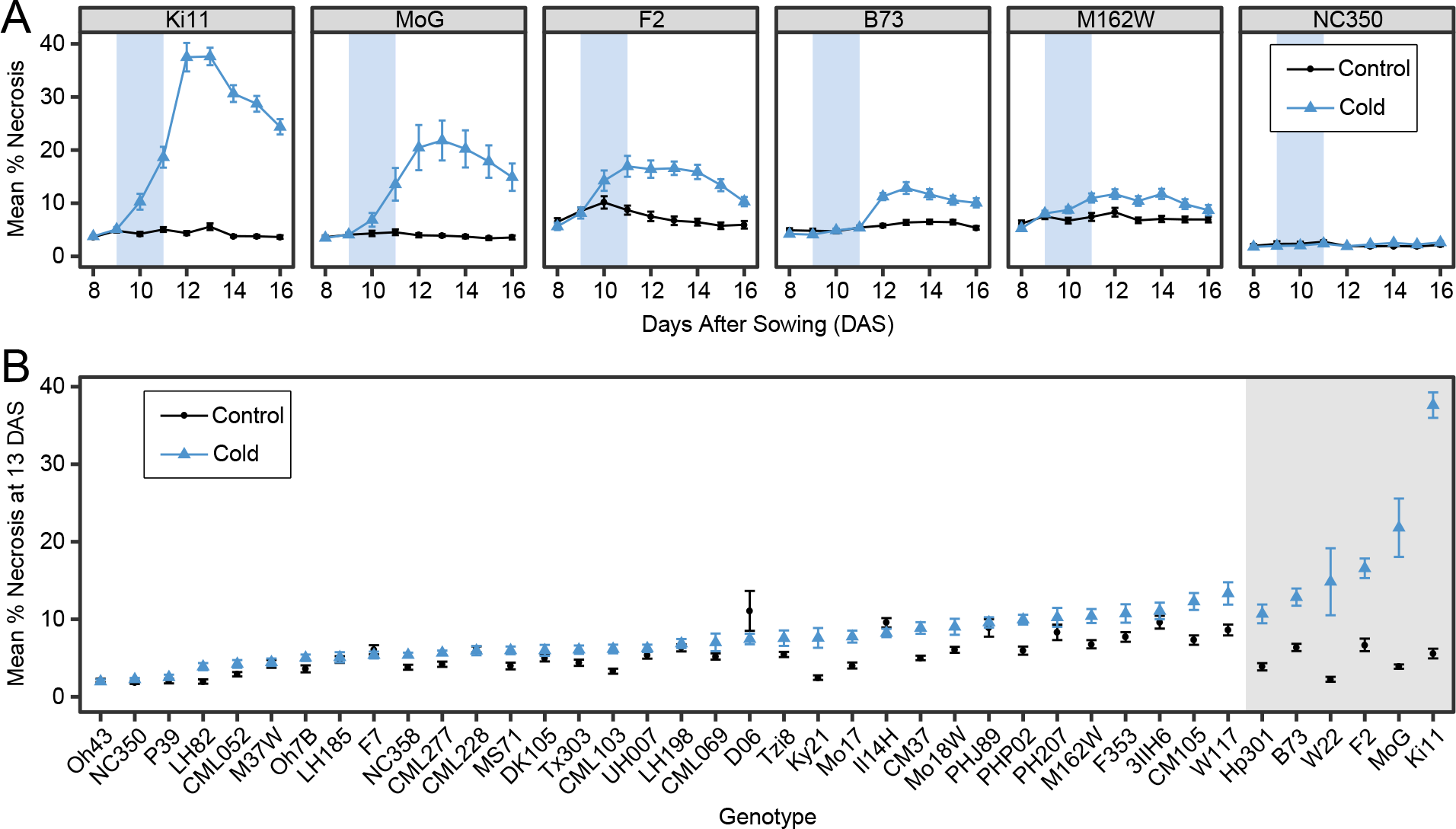
Percent necrosis over time. A) Mean percent necrosis for each time point for 6 maize genotypes. Error bars represent standard error of the mean. B) Mean percent necrosis on day 13 for 40 maize genotypes. Error bars represent standard error of the mean. Data shaded with a gray box had an adjusted p-value of ≤ 0.05 in a one-way ANOVA and post-hoc Tukey test.

Among the genotypes surveyed, we observed substantial variation for this response (Figure 5A, Supplemental Figure 2). Some genotypes, such as Oh43 and NC350, did not exhibit any changes in the proportion of necrotic tissue following a cold stress (Figure 5A). For other genotypes, such as MoG or Ki11, a significant portion of the plant exhibited necrosis. A comparison among these genotypes also revealed variability in the level of necrotic tissue present within the control plants. This resulted from healthy portions of the lower stem having color values with a high probability of being classified as necrosis. This occurred at different frequencies among genotypes. To control for this, we focused on comparing the amount of necrotic tissue in control plants compared to cold stressed plants within each genotype. We also noted variability in the total percent of necrotic tissue during our time series. The percent necrotic value often peaked at 13 DAS followed by a gradual decline. This was a result of new growth of healthy/green tissue following the cold stress, resulting in the overall percent necrotic area decreasing. The percentage of plant area that exhibited necrosis was finite and underwent a predictable sequence of color changes. Therefore, a time-course analysis is unnecessary for this phenotype within our experiments.

Accordingly, the percent necrotic tissue was assessed for all 40 genotypes at 13 DAS (Figure 5B). Seven genotypes had a significant increase in tissue classified as necrosis relative to the controls. For many of the other genotypes, the cold treatment group never accumulated an amount of necrotic tissue greater than the amount of tissue misclassified as necrotic within the control group. It is worth noting that all genotypes survived the cold stress and continued growth. Even the most severe necrosis responses only resulted in the loss of healthy tissue for two or three leaves.

### Impact of cold stress on plant morphology across genotypes

Genotypes were compared on a single day for quantification of necrotic tissue, however the morphological traits changed at different rates among genotypes, and therefore the entire time course of data collected was utilized (Supplemental Figures 3-7). For example, plant area did not increase during the cold treatment for any genotype (Supplemental Figure 3). Yet, the rate at which area increased following the cold stress period was faster for some genotypes compared to others (Figure 6A, Supplemental Figure 3). Growth rates across traits are more similar within genotypes than among genotypes. The set of traits that are most affected by stress vary across genotypes. Values for all analyzed morphological traits did not appear to increase during the cold treatment for any genotype but recovery rates following the stress varied. Because genotypes are not growing during the cold stress period, they are delayed in development compared to plants grown under control conditions. Because of the cold-induced delay in growth, a more equivalent comparison than comparing control and cold-stressed data at the same time point was to compare cold-stressed data to control data at a time point two days earlier. Therefore, we chose to compare measurements for morphological traits with a time-shifted approach, comparing the cold-stress plant measurements to the control plant measurements collected 2 days prior for each genotype (Figure 6B). This allowed comparisons of control and cold stressed plants at more similar developmental stages. Additionally, to minimize the effects of individual plant size on comparisons, we calculated growth rates during two intervals among the time-shifted data (Figure 6C). Within each interval, we calculated the log_2_ fold change in growth rates between treatments for each genotype for each trait.

**Figure 6.**
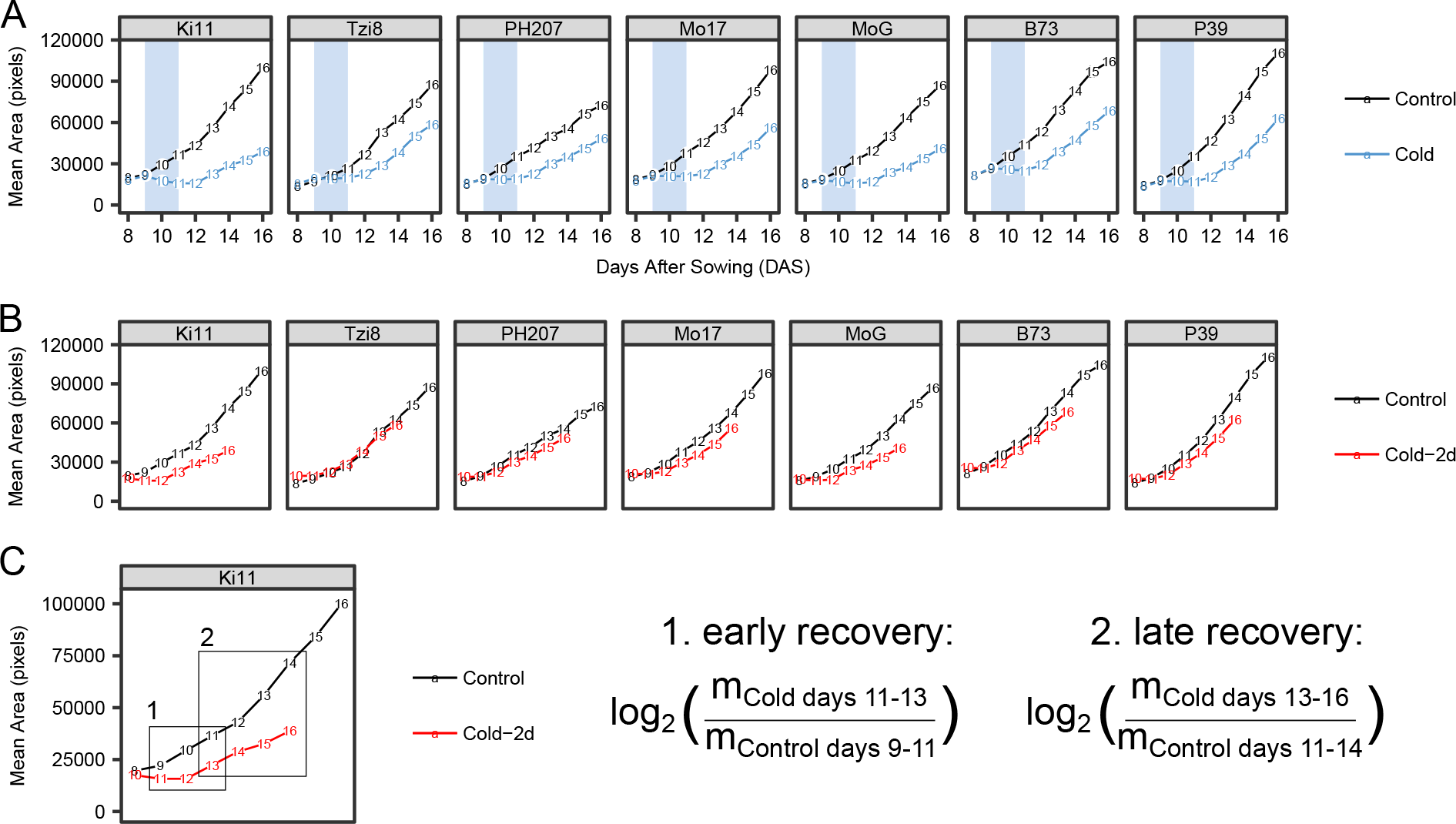
Morphological traits over time and calculations for clustering. A) Mean area for each time point for 7 maize genotypes used in this study. B) Data presented in panel A with 2 days shift back in time for cold-treated samples. C) Selection of time intervals for comparing growth rates during equivalent developmental stages of control and cold-treated plants.

### Clustering genotypes based on leaf necrosis and morphology phenotypes

Because of intrinsic morphological differences among genotypes, growth rates manifest in different patterns across traits. Therefore, a phenotype fingerprint, or phingerprint, captures the unique aspects of cold response in each genotype. The time-shifted and normalized data from plant morphology phenotypes and the percent necrosis data from day 13 was used to cluster genotypes using hierarchical clustering (Figure 7). This resulted in three clusters. Overall patterns of growth inhibition were fairly similar across all genotypes. However, subtle differences among degrees of fold change and the pattern of changes across different morphological traits helped define genotypes into different clusters. One cluster was characterized by more subtle changes in growth rates compared to controls of morphological traits over the two defined intervals. A second cluster was dominated by higher degrees of necrosis than controls on day 13. A third cluster was characterized by moderate levels of necrosis on day 13 and a smaller fold change in growth rate of height compared to controls. Our initial hypothesis was that genotypes may cluster together based on some attributes, such as population group, market class, kernel type, or latitude. This did not appear to be the case. Latitude did not have a significant correlation with any of the log_2_ fold changes in growth rates for any traits during either interval. Therefore, cold sensitivity during early development was likely not a target of breeding programs for these genotypes.

**Figure 7.**
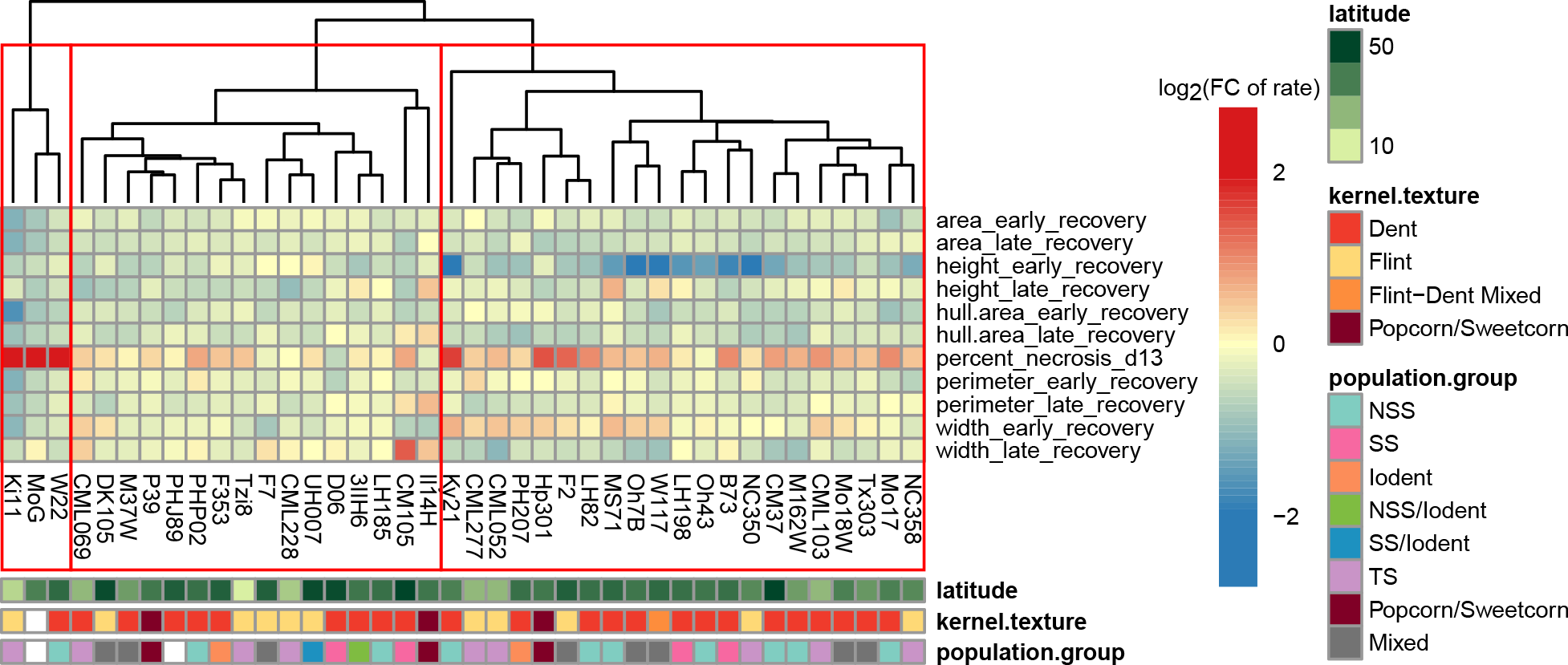
Phingerprints classify maize inbreds into three clusters. Each cell represents the log_2_ fold change in growth rate between control and cold-treated plants for the trait and interval indicated for each row.

## Discussion

Improving cold tolerance in maize remains a challenge, despite nearly 100 years of studies. Phenotyping approaches can enable new insights into cold stress responses in plants. This study successfully used image-based phenotyping methods to characterize how maize seedlings respond and recover from a cold stress event. Our approach allowed comparisons of multiple genotypes over time and for quantitative measurements of cold stress responses, such as area, height, width, and development of leaf necrosis. The quantification of these traits from images facilitated nondestructive measurements to be made over time and enabled our analysis of recovery rates. The use of hierarchical clustering allowed for a methodical approach to compare the ability of genotypes to recover from cold stress.

As with most image-based time course phenotyping methods, there were several limitations to our approach. The patterns of clustering were influenced by the choice of time intervals and included traits. Every inbred in our study is sensitive to cold, so differences among that sensitivity can be subtle. The quantification of the leaf necrosis phenotype provided a straightforward definition of the most cold sensitive genotypes within our assay, however the morphological traits captured subtle variations in the diverse patterns of cold response. Additionally, although our assay recapitulates phenotypes observed in field-grown plants that experience cold stress (chlorotic bands on leaves and leaf necrosis), caution must be made when extrapolating controlled-condition studies to field conditions. Finally, the performance of inbreds does not always predict combining ability for hybrids, so future studies could include analysis of heterosis among various hybrids for recovery from cold stress.

Many studies suggested flint genotypes are more cold tolerant than dent genotypes across a number of different cold assays and developmental stages (Bhosale et al., 2007; Riva-Roveda et al., 2016). With our approach, flint and dent genotypes did not classify into separate clusters, although dent genotypes did appear to have more severe necrosis than flint genotypes. Ki11, a flint genotype, exhibits the highest percentage of necrosis of all genotypes in our study. Other examples violating the assumption that dent genotypes are cold sensitive exist, such as a favorable allele for cold tolerance being identified in a European dent line (Strigens et al., 2013). Additionally, in a large study comparing cold tolerance of flint and dent varieties of European origin, genotypes with a high degree of cold tolerance were found among both dent and flint groups (Revilla et al., 2014). The same study found no strong pattern among cold tolerance and latitude of geographical origin, which is also consistent with our results.

Assigning a latitude and other attributes, such as kernel texture and population group, to maize inbreds can be a difficult task. Many genotypes have complex pedigrees, making distinct category assignments inaccurate or not representative of breeding history. For example, Mo17 was developed in the state of Missouri in the United States; however, lines used to generate Mo17 were developed in both Illinois and Connecticut (Andorf et al., 2016). This example illustrates how assigned geographical regions of maize inbreds may not be straightforward and could help explain the lack of correlation between our measures of cold sensitivity and latitude. Alternatively, it is possible that different breeders utilized different practices in terms of planting date. Breeders that preferred to plant earlier in the season likely imposed some selection for early season cold tolerance while breeders that planted later in the season likely did not impose this selective pressure.

Few studies have analyzed responses to cold stress in maize over time. One recent study that performed time-series analysis of leaf elongation rates under a cold stress used manual methods to collect this data (Riva-Roveda et al., 2016). Measurements collected using image-based phenotyping can greatly reduce the amount of time and labor needed to investigate how traits change over time, as well as archive plant morphology for future studies. Our approach enabled the simultaneous collection of multiple morphological and color-based measurements at nine time points for thousands of individual plants. Having daily imaging and quick trait extraction pipelines allowed us to collect a large amount of phenotypic data on many plants. Using these techniques, we recognized the delay in growth rates the stress caused, and therefore we compared treatment groups at more equivalent developmental stages. This analysis would not have been possible if we had made measurements at fewer time points.

Our goal was to provide a basis for designing genetic mapping studies to work towards maximizing the ability of maize to withstand and recover from early season cold stress events. Additional future studies could include a cold acclimation period and analyzing whether this impacts recovery rates, leaf necrosis, and the clustering of genotypes. Image-based phenotyping methods can provide more in-depth analysis of cold stress responses in maize and may help tease apart the narrow genetic variance for cold tolerance in this important crop species.

## Acknowledgments

We thank Veronica Swanson, Meg Gerold, Hayden Christensen, Kjell Sandstrom, Danielle Sorenson, and Shale Demuth for technical assistance in data collection. This work was supported by the National Science Foundation Plant Genome Award IOS-1444456.

## Author Contributions

N.M.S., C.D.H., and T.A.E. designed the research; T.A.E., S.S.D., and J.O. performed research; S.T.C., N.D.M., C.D.H. contributed new computational tools; T.A.E. analyzed data; T.A.E., N.M.S., and C.D.H. wrote the paper. All authors commented on and approved the final manuscript.

## Supplemental Data

Supplemental Table 1. Information about each genotype.

Supplemental Table 2. Sample size information.

Supplemental Figure 1. Image output from PlantCV pipeline. A) Merged image indicates pixels categorized as healthy (blue) or necrotic (red). B) Final plant binary mask used for shape and color analysis. C) Output from PlantCV analyze_object function. D) Output from PlantCV analyze_color function false colored using the “value” channel from HSV colorspace.

Supplemental Figure 2. Mean values for percent necrosis at each time point for 40 maize inbred genotypes. Error bars represent standard error of the mean.

Supplemental Figure 3. Mean values for plant area at each time point for 40 maize inbred genotypes. Error bars represent standard error of the mean.

Supplemental Figure 4. Mean values for plant height at each time point for 40 maize inbred genotypes. Error bars represent standard error of the mean.

Supplemental Figure 5. Mean values for plant width at each time point for 40 maize inbred genotypes. Error bars represent standard error of the mean.

Supplemental Figure 6. Mean values for plant perimeter at each time point for 40 maize inbred genotypes. Error bars represent standard error of the mean.

Supplemental Figure 7. Mean values for plant hull area at each time point for 40 maize inbred genotypes. Error bars represent standard error of the mean.

